# A phospholipase assay screen identifies synergistic inhibitors of the *P. aeruginosa* toxin ExoU

**DOI:** 10.1101/2022.02.21.481271

**Authors:** Daniel M. Foulkes, Keri McLean, Anne Hermann, James Johnson, Craig Winstanley, Neil Berry, David G. Fernig, Stephen B. Kaye

## Abstract

The opportunistic pathogen *Pseudomonas aeruginosa* is a leading cause of disability and mortality worldwide and the World Health Organisation has listed it with the highest priority for the need of new therapies. *P. aeruginosa* strains that express ExoU are implicated in the worst clinical outcomes. ExoU is phospholipase that is secreted by *P. aeruginosa* directly into the cytoplasm of target host cells, where its catalytic activity, directed towards plasma membranes, causes rapid cell lysis. Inhibition of ExoU may be a novel strategy to combat acutely cytotoxic ExoU expressing *P. aeruginosa* infections. Using an *in vitro* phospholipase assay, we performed a high throughput screen to identify compounds that might be repurposed as therapeutic ExoU inhibitors. We discovered a panel of compounds that appeared to inhibit ExoU through distinct mechanisms. Compound C prevented ExoU membrane localisation in HEK293T cells and caused colocalization with lysosomes, whereas compound D prevented PIP_2_ dependent oligomerisation of ExoU *in vitro* suggestive of synergistic action. Indeed, the concentrations required by compounds C and D to inhibit *in vitro* ExoU catalytic activity, when used in combination, was in the nanomolar region. In corneal scratch and infection assays, these compounds reduced ExoU mediated cytotoxicity, as assessed by fluorescence microscopy and lactate dehydrogenase release assays.

## Introduction

The opportunistic pathogen *P. aeruginosa* is a major cause of nosocomial infection, associated with a wide range of debilitating and fatal diseases [1–3]. It is a leading cause of intensive care unit-acquired pneumonia (ICUAP) [4], and is the second most frequent colonising bacteria in patients with COVID19 [5, 6]. It is also the primary causative agent of bacterial keratitis [7]. As a pathogen of current major concern, the World Health Organisation (WHO) has listed carbapenem-resistant *P. aeruginosa* (CRPA) with the highest priority for the development of new treatments [8].

Pathogenic *P. aeruginosa* strains use the type III secretion system (T3SS) to inject exotoxins directly into the cytoplasm of compromised host epithelia [9]. The T3SS has been identified as a principal virulence determinant for poor clinical outcomes in pneumonia, sepsis, keratitis, and otitis externa [2–4, 10, 11]. Of the four effector proteins secreted by the T3SS, ExoS, ExoT, ExoU and ExoY, the most cytotoxic is ExoU. In *P. aeruginosa* infections where ExoU is expressed, disease severity is increased with poorer clinical outcomes [2, 4, 12, 13].

ExoU is a potent phospholipase that is secreted directly into the target host cell cytoplasm via the *P. aeruginosa* T3SS [9]. Although the mechanisms of ExoU activation are not fully understood, it is established that ExoU binds to ubiquitin and phosphatidylinositol 4,5-bisphosphate (PIP_2_) to become activated [14, 15]. PIP_2_ induces ExoU oligomerisation, by which its ubiquitin-dependent catalytic activity is greatly enhanced [15]. In mammalian cells, binding of PIP_2_ causes ExoU localisation to the plasma membrane, via its C-terminal 4-helical bundle domain, where its catalytic activity induces cellular lysis [16, 17]. ExoU catalytic activity is directed towards phospholipids at the sn-2 position, and results in arachidonic acid release, which in turn stimulates NF-κB and MAPK signalling [18–20]. Increased expression of IL-8 and keratinocyte chemoattractant (KC) then causes acute localised inflammation, leading to infiltration of neutrophils and further tissue damage [18, 19].

Inhibitors of ExoU might reduce acute cell lysis, tissue damage and inflammation associated with ExoU toxicity and provide a more substantial window for immune response or antibiotic action [21]. ExoU undergoes multiple activation steps including interaction with ubiquitin, DNAJC5 dependent host trafficking [22] and localisation to plasma membranes, which might be targetable with small molecules. In this study, we performed an *in vitro* phospholipase assay screen in order to identify novel ExoU inhibitors, interrogate their mechanisms of inhibition and test their efficacy in a corneal cell infection model.

## Materials and methods

### Materials

The pOPINF *E. coli* expression vector was purchased from Addgene. The pEGFP-C3 vector, in which ExoU was cloned for expression of GFP-tagged ExoU in human cells, was a gift from Dr. Michael Batie (University of Liverpool, UK). Arachidonoyl thio-PC was purchased from Cayman Chemical (Michigan, USA). PIP_2_ was purchased from Avanti Polar Lipids (Alabama, USA). Bovine ubiquitin and 5,5’-dithiobis-(2-nitrobenzoic acid) (DNTB) were purchased from Sigma-Aldrich. ExoU screening compounds were provided by LifeArc (UK) and Professor Patrick Eyers (University of Liverpool, UK). The α-GFP and α-tubulin antibodies were purchased from Sigma Aldrich. Live/Dead assay reagents were purchased from Invitrogen. LDH assay reagents and propidium iodide were purchased from Thermofisher. The *P. aeruginosa* strain PA103 was a gift from Professor Dara Frank (Medical College of Wisconsin).

### Recombinant protein production

Full-length ExoU, expressed in pOPINF, was purified from C43(DE3) *E. coli* as previously described [23]. Briefly, transformed C43(DE3) bacteria were grown in Terrific broth (Melford) supplemented with ampicillin (100 μg/ml) and grown to an optical density (OD600) of 0.8, at which point 0.4 mM isopropyl-β-d-thiogalactopyranoside (IPTG) was added to induce His-ExoU expression overnight at 18°C. The bacterial lysis buffer contained 20 mM Tris–HCl pH 8.2, 300 mM NaCl, 0.1% (v/v) Triton-X-100, 10 mM imidazole, 1 mM DTT, 10% (v/v) glycerol and a cOmplete protease inhibitor cocktail tablet (Roche) and 100 μM Phenylmethylsulfonyl fluoride (PMSF). ExoU was purified by an initial affinity step (immobilised nickel affinity chromatography) followed by size-exclusion chromatography (SEC) (16/600 Superdex 200, GE healthcare) in 20 mM Tris–HCl pH 8.2, 100 mM NaCl and 10% (v/v) glycerol. Recombinant ExoU was frozen in liquid nitrogen and stored at −80°C.

### ExoU phospholipase assay compound screen

Recombinant ExoU sn-2 directed phospholipase activity was detected using an adapted Caymen chemical cPLA2 assay kit in a 384-well plate format, as previously described [23]. A master mix contained 1mM ATPC, 1 μM PIP_2_, 25 μM mono ubiquitin, 1.25 mM 5,5-dithio-bis-(2-nitrobenzoic acid) (DTNB) and 100 nM ExoU. The master mix was dispensed into 384-well plates at a volume of 10 μL per well. An Echo acoustic liquid handling robot was then used to dispense 0.2 μL of screening compound to each well. The final concentration of compound for each condition 10 μM. ExoU activity was detected as percentage to that of DMSO controls, 4 hours after the start of the assay.

### ExoU oligomerisation assay

Compounds or DMSO (2% v/v) were added to 1 μg of ExoU with 2.5 μM of PIP_2_ in 20 mM Tris–HCl pH 8.2, 100 mM NaCl and 10% (v/v) glycerol and incubated on ice for either 30 min or 16 h. Sample buffer (0.5 M Tris–HCl pH 6.8, 25% v/v glycerol and 0.25% v/v bromophenol blue) was added to each sample prior to resolution on 12 % Mini-PROTEAN TGX Gels (Bio-Rad) by polyacrylamide gel electrophoresis (PAGE) with sodium dodecyl sulphate (SDS) absent from the running buffer. After PAGE, proteins were transferred to nitrocellulose membranes (Bio-Rad) which were blocked in Tris-buffered saline + 0.1% (v/v) Tween 20 (TBS-T) in 5% (w/v) milk (pH 7.4) followed by incubation with an anti 6xhistidine antibody (Bio-Rad) overnight. Proteins were detected using secondary HRP-conjugated α-mouse antibody and enhanced chemiluminesence reagent (Bio-Rad).

### Human cell transfections

Adherent parental Flp-In T-REx-HeLa (Invitrogen) and HEK293T were cultured in Dulbecco’s modified Eagle medium (DMEM) supplemented with 4 mM l-glutamine, 10% (v/v) foetal bovine serum (FBS), Penicillin and Streptomycin (Gibco), 4 μg/ml of Blasticidin (Melford) and Zeocin 50 μg/ml (Invitrogen). Cells (2.2 × 106 cells in 10 cm dishes and 0.5 × 106 cells for 6-well plates) were seeded 24 h prior to transfection. Cells were transfected with either pcDNA5/FRT/TO plasmid encoding Flag-tagged WT ExoU, or pEGFP-C3 to express GFP-tagged S142A ExoU, using polyethylenimine (PEI) at a ratio of 3:1 PEI:cDNA.

### Trypan blue assay

Transfected HeLa cells were collected 8 h after transfection and tetracycline (TET) induced ExoU expression, by combining the culture medium (to obtain suspended cells) and adherent cells released by trypsinisation. The resulting HeLa cell and medium suspension was mixed in a 1:1 ratio with trypan blue reagent (Thermo Scientific) and a Countess II Automated Cell Counter (Thermo Scientific) was employed to detect the percentage of lysed cells

### Propidium iodide uptake

Transfected HeLa cells (including suspended cells) were collected by trypsinisation 8 h after induction of ExoU expression in presence of the indicated compound. Samples were stained with propidium iodide (PI) and diluted with PBS so that samples contained less than 500 cells/μl for 10 000 events per run. After gating to select whole cells, employing a BD Accuri C6 flow cytometer, the total cell numbers were evaluated with forward scatter and PI-stained cells were detected by using an appropriate laser for fluorescence. Results were given as relative florescence units for PI uptake.

### Western blotting

Western blotting was used to detect total GFP-S142A ExoU in transfected HEK293T cells. whole-cell lysates were generated using a modified RIPA buffer (50 mM Tris–HCl pH 7.4, 1% (v/v) NP-40, 0.1% (v/v) SDS, 100 mM NaCl, 1 mM DTT, 10% (v/v) glycerol, a complete protease inhibitor cocktail tablet (Roche). After resuspension in lysis buffer, HEK293T cells were briefly sonicated and centrifuged at 16 000g prior to quantification of protein concentration using Bradford reagent. Samples were boiled in sample buffer (50 mM Tris–Cl pH 6.8, 1% (w/v) SDS, 10% (v/v) glycerol, 0.01% (w/v) bromophenol blue, and 10 mM DTT) and 40 μg of total protein, for each sample, resolved by SDS–PAGE prior to transfer onto nitrocellulose membranes (Bio-Rad). Membranes were blocked in Tris-buffered saline (pH 7.4) with 0.1% (v/v) Tween 20 (TBS-T) in 5% (w/v) milk followed by incubation with primary (α-GFP) and secondary (α-mouse-HRP) antibodies. Proteins were detected using enhanced chemiluminesence reagent (Bio-Rad).

### GFP-ExoU localisation experiments

HEK 293T cells were seeded on 4 compartment glass-bottom imaging dishes (Ibidi) and transfected with pEGFP-C3 to express GFP-tagged S142A ExoU and RFP-Lamp1 (ThermoFisher), when 70% confluent. 2 hours after transfection, cells were incubated with DMSO (0.1% v/v) or compound C. Intracellular localisation of treated or untreated ExoU was monitored live and longitudinally for 24h by confocal microscopy (Zeiss LSM780) at 37 °C and 5% CO_2_. Colocalisation, as well as colocalisation coefficients for ExoU and lysosomes, were analysed with ImageJ.

### HCE-T scratch and infection assay

HCE-T cells were seeded in 6-well plates and grown until a fully confluent monolayer had formed. A 10 μl pipette tip was used to make a scratch across the diameter of the wells. PA103 that had previously been expanded in LB culture to an OD_600_ of 0.8 were added to the scratched HCT cells at an MOI of 10. Compounds or DMSO (0.1 % v/v) were also added and infection was allowed to take place for 6 hours, after which Live/Dead fluorescence microscopy and LDH assay analyses were performed.

### Live/Dead fluorescence microscopy

Scratched and infected HCE-T cells, in the presence of indicated compounds, were analysed by florescent microscopy, employing Live/Dead staining (Invitrogen), according to the manufacturer’s instructions. After 6 hours PA103 infection, HCE-T cell culture medium was removed and cells were washed with 1 ml of PBS three times. Fresh medium, containing 5 μM of both calcein (Ex/Em 494/517 nm) and ethidium homodimer-1 (Ex/Em 528/617 nm), was then added and images of the scratched HCT cells were obtained on either a Nikon Eclipse TiE.

### LDH assays

Lactate dehydrogenase (LDH) release was measured using the Pierce LDH Cytotoxicity Assay Kit (Thermo Scientific) according to the manufacturer’s instructions. Culture medium (50 μl) from transfected HeLa or infected HCE-T cells were assayed after 6 hours, in the presence of specified compounds (0.1% v/v DMSO). The absorbance of the negative controls (untransfected/uninfected cells) was subtracted to yield the final absorbance values

## Results

### Discovery and biochemical analysis of ExoU inhibitors

ExoU with an N-terminal histidine tag was purified to homogeneity from *E. coli* by immobilised metal affinity chromatography (IMAC) and then by size-exclusion chromatography (SEC) (Figure 1A). The phospholipase activity of ExoU was measured by adaptation of the Cayman chemical PLA2 phospholipase assay kit, so that analysis was compatible with a 384-well plate format for a screen of 6,500 compounds (Figure 1B). Arachidonoyl Thio-PC substrate hydrolysis by 100 nM of recombinant ExoU in 2% (v/v) DMSO was assessed in the presence of 10 μM of compound, in a 4-hour end point assay. The most potent inhibitors of ExoU that were discovered we have named compound C (IC_50_ = 1.7 ± 0.5), compound D (IC_50_ = 16.3 ± 3.4), compound E (IC_50_ = 37.7 ± 5.4) and compound F (IC_50_ = 29.6 ± 4.3).

**Figure 1:**
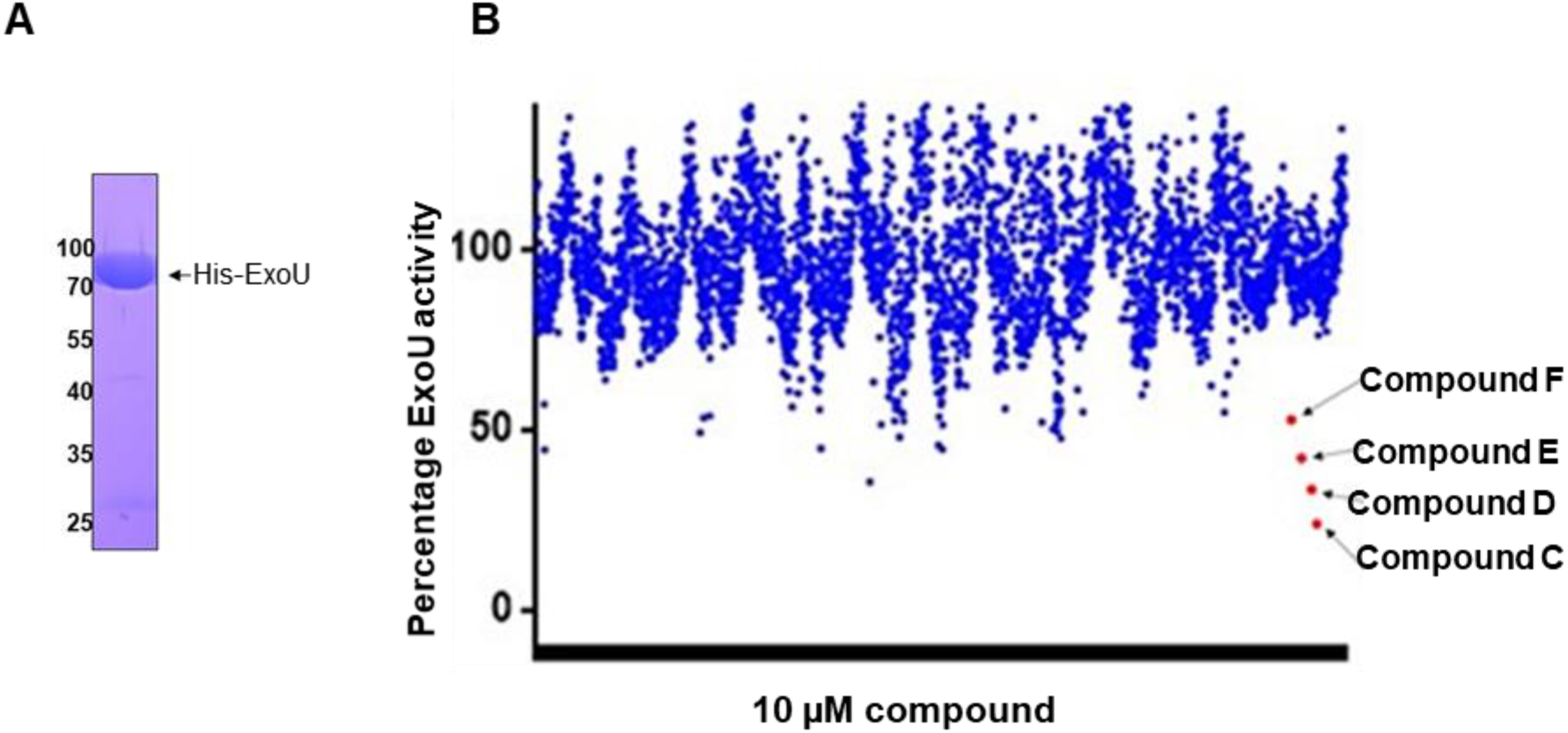
An *in vitro* phospholipase assay screen to identify new ExoU inhibitors: **(A)** SDS-PAGE analysis of recombinant His-tagged ExoU, purified from *E. coli,* by immobilised metal affinity chromatography (IMAC) and size exclusion chromatography (SEC). **(B)** A screen of 6500 small molecules using an *in vitro* phospholipase assay to identify compounds that could inhibit ExoU activity. In each condition, 100 nM of ExoU was assayed in the presence of 10 μM of compound, with ubiquitin and PIP_2_ present as activators of ExoU. After 4 hours, substrate cleavage by ExoU was detected as a function of absorbance at 405 nm and converted to percentage activity with reference to DMSO controls.

*In vitro* ExoU forms oligomers in the presence of PIP_2_, which in turn greatly enhances ExoU catalytic activity. Using native-PAGE and immunoblotting we found that compound D, but not compound C, E or F, was able to prevent ExoU oligomerisation *in vitro* (Figure 2). SpcU, the cognate inhibitory chaperone of ExoU, also prevented ExoU oligomerisation, serving as a positive control (Figure 2).

**Figure 2:**
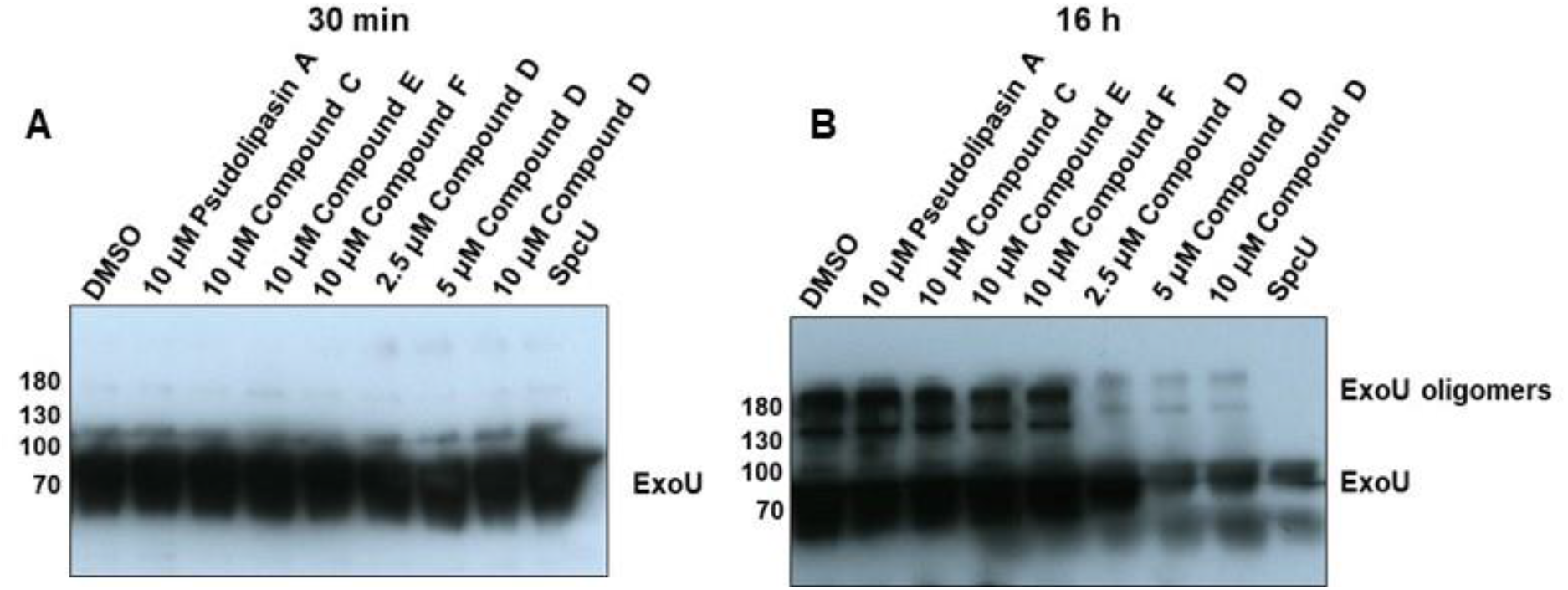
Compound D prevents ExoU oligomerisation. Native-PAGE and immunoblotting analysis of recombinant ExoU in the presence of compounds and SpcU to detect ExoU oligomerisation. 1 μg of ExoU was incubated for (A) 30 minutes or (B) 16 hours with 2.5 μM of PIP_2_ in the presence of indicated compound or the cognate ExoU chaperone SpcU, prior to native-PAGE and immunoblot analysis.

We hypothesised that the mechanism of inhibition of ExoU by compound D was distinct to that of compound C, E and F, and therefore might be used in combination to further increase ExoU inhibitory activities. Alone, compounds C and D inhibited ExoU phospholipase activity, with IC_50_ values of 1.7 ± 0.5 μM and 16.3 ± 3.4 μM, respectively (Figure 3A). However, the inhibitory activity for these compounds, when used in combination, is markedly improved to an IC_50_ of 310 ± 16 nM (Figure 3B).

**Figure 3:**
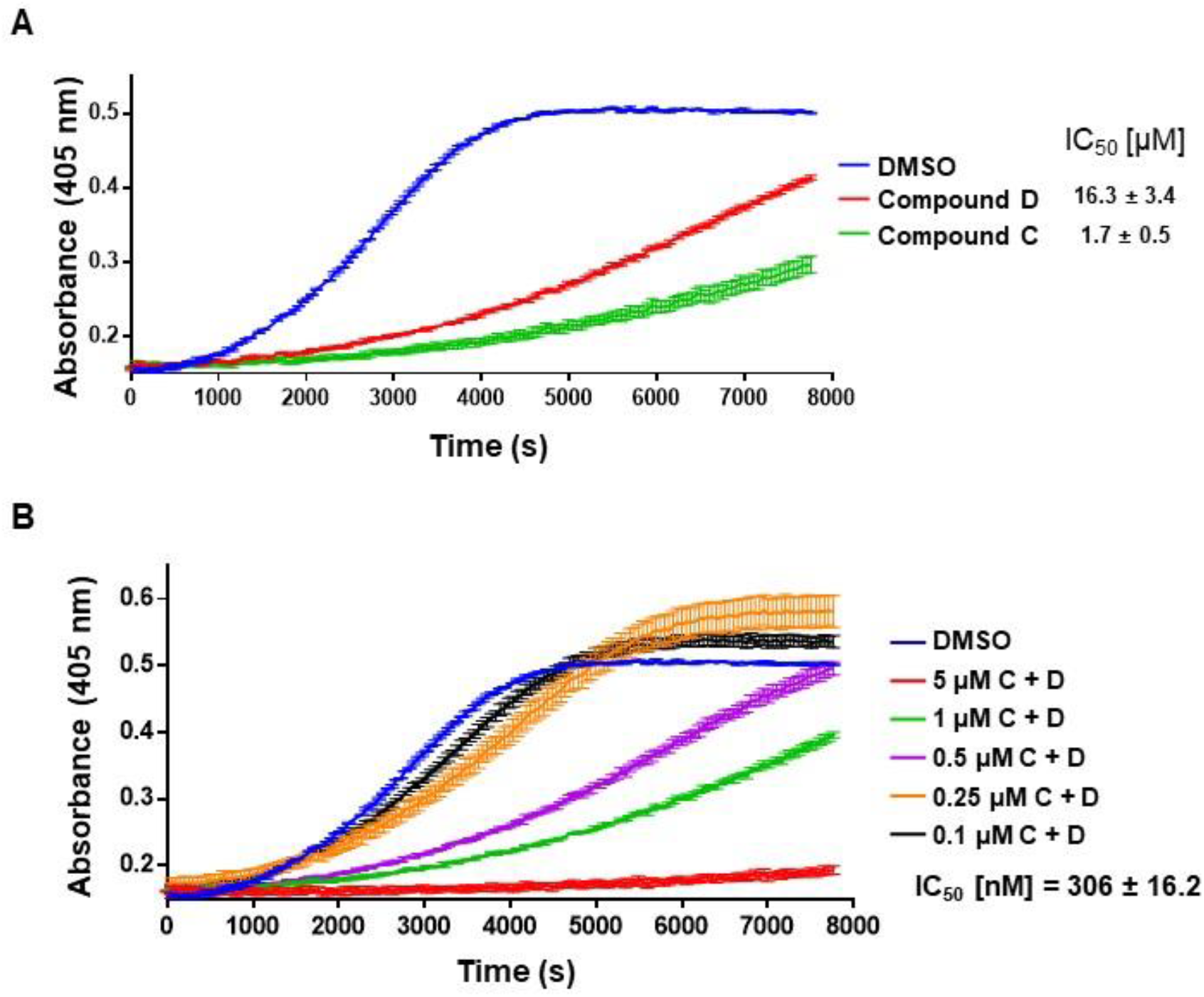
Compounds C and D synergise to inhibit ExoU phospholipase activity *in vitro.* (A) The hydrolysis of arachidonoyl Thio-PC substrate by ExoU was assessed in the presence of 10 μM of the indicated compound. To each reaction, ubiquitin and PIP_2_ were added in order to allow induction of ExoU phospholipase activity. (B) Dose dependent inhibition of ExoU phospholipase by the combination of compound C and D, present at equimolar concentrations.

### Analysis of ExoU inhibitors in a HeLa transfection system

Human cells transfected to express WT ExoU undergo membrane blebbing and rapid cell lysis [24]. Using a previously established protocol [23], we sought to determine whether or not compounds C, D, E or F could protect transfected HeLa cells from ExoU mediated cell lysis (Figure 4). Brightfield microscopy was employed to observe morphology and quantity of adherent HeLa cells after FLAG-tagged ExoU expression, with and without compounds present in HeLa cells (Figure 4A). The percentage LDH release (Figure 4B) and propidium iodide (PI) uptake (Figure 4C) were used to quantity the extent of cell lysis caused by ExoU, with and without compounds present. For HeLa cells transfected to express FLAG-ExoU, there was a greater number of rounded cells and fewer cells remained adhered to the well bottom, compared to vehicle transfected cells. Expression of ExoU was accompanied by an increase in percentage LDH release (Figure 4B) and propidium iodide uptake (Figure 4C). At 10 μM, compounds C, D, E and F all appeared to mitigate the effects of ExoU expression, with fewer visibly rounded HeLa cells and decreases (similar for all compounds) in LDH release and PI uptake.

**Figure 4:**
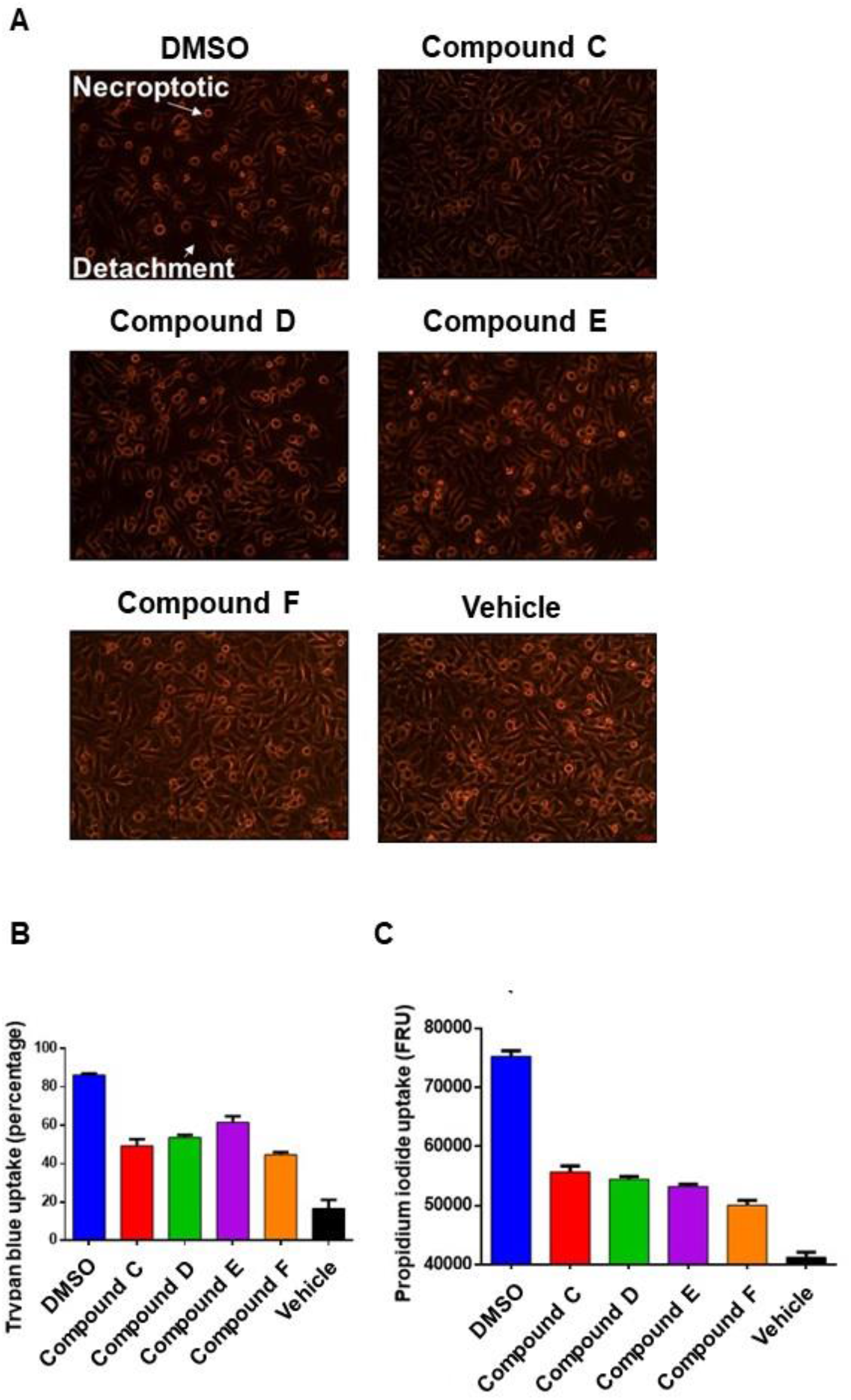
Compounds reduce toxicity in ExoU expressing transfected HeLa cells. Flp-In T-REx parental HeLa cells transfected with pcDNA5/FRT/TO, encoding WT ExoU, to allow for tetracycline inducible expression of ExoU. After induction of ExoU expression, 10 μM of inhibitor was added for 8 hours. (A) Brightfield microscopy, (B) trypan blue uptake and (C) propidium iodide uptake analyses were used to assess the protective effects of compounds from ExoU toxicity.

### ExoU localisation influenced by Compound C

In a previous study, we determined that established ExoU inhibitors, Pseudolipasin A, compound A and compound B, caused a reduction in the total amount of ExoU observed in transfected human cells. We sought to determine how the compounds identified in the present screen influenced ExoU stability and localisation in transfected HEK293T cells. As wild-type ExoU causes rapid lysis when expressed in human cells, HEK293T cells were transfected with pEGFP-C3 to express a GFP-tagged catalytically inactive (S142A) ExoU mutant (Figure 5). Compound C caused a dose-dependent reduction in the total quantity of GFP-S142A ExoU, detectable by western blotting, in the lysates of transfected HEK293T cells. Employing fluorescence microscopy, we observed that without compound present (DMSO 0.1% v/v), GFP-ExoU localised almost exclusively to plasma membrane (Figure 5B). However, in the presence of compound C, GFP-S142A ExoU was observed to be punctate throughout the cytoplasm (Figure 5B). Our previous study indicated that ExoU was not degraded by proteasomes, so we hypothesised that certain compounds might cause lysosomal degradation of ExoU. To this end, we co-transfected HEK293T cells to express GFP-S142A ExoU and RFP-tagged lysosomal associated protein 1 (RFP-LAMP1) (Figure 5C). We found that GFP-S142A ExoU colocalised RFP-LAMP1 (Figure 5C), which might indicate that, in the presence of compound C, ExoU is targeted for lysosomal destruction.

**Figure 5:**
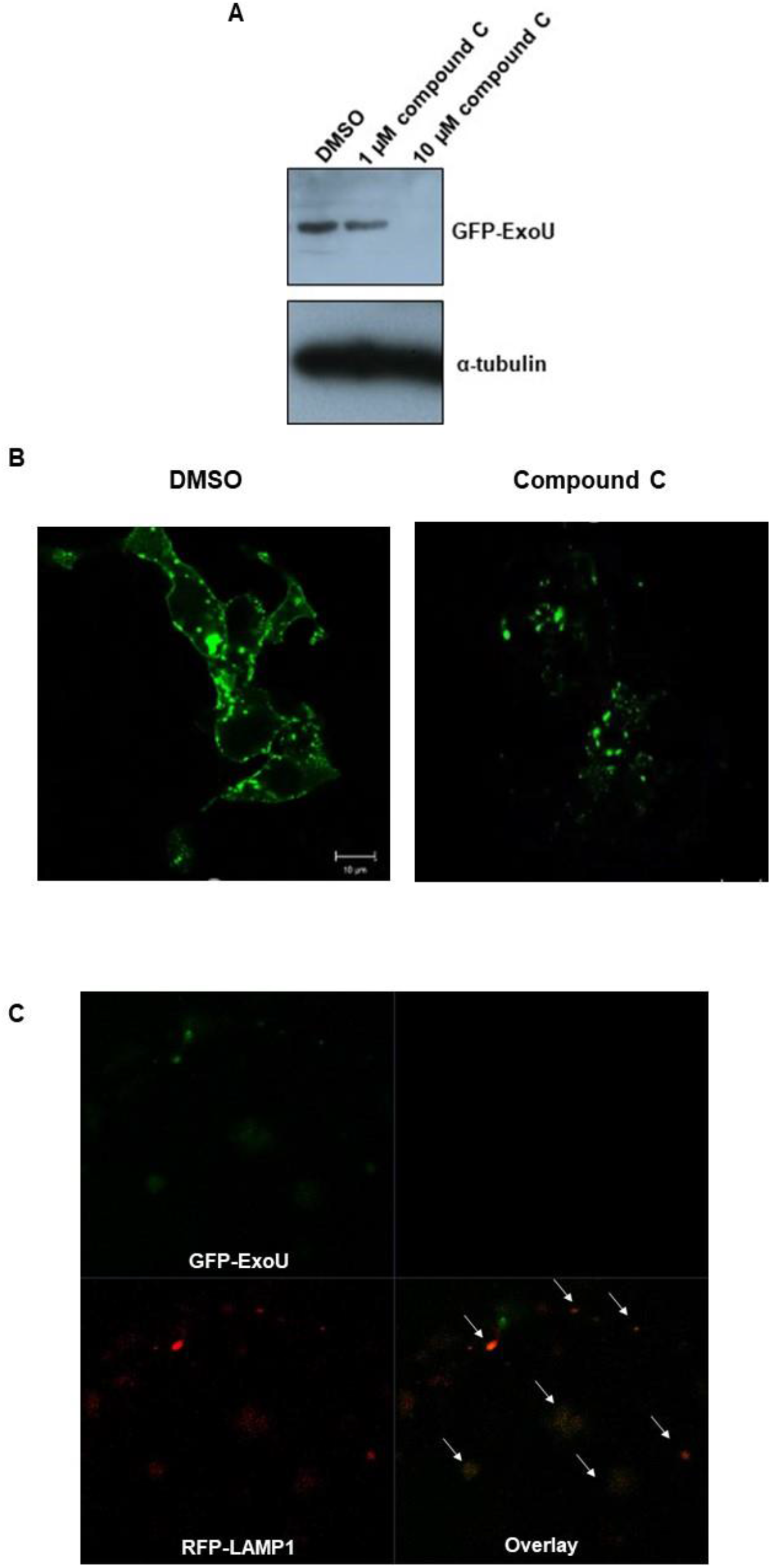
Compound C induces lysosome localisation and degradation of GFP-ExoU in HEK293T cells. (A) HEK293T cells were transfected with pEGFP-C3, encoding S142A ExoU, in the presence of compound C for 16 hours. Western blotting of whole cell lysates was used to detect abundance of GFP-ExoU in transfected HEK293T cells, using an α-GFP antibody. (B) Transfected HEK293T cells were incubated ± 1 μM compound C for 16 hours and analysed by fluorescence microscopy. (C) HEK293T cells were cotransfected (± 1 μM compound C) to express GFP-ExoU and RFP-tagged lysosome associated protein 1 (LAMP1). Colocalization of GFP-ExoU RFP-LAMP1 was assessed by fluorescence microscopy.

### Analysis of compounds in a *P. aeruginosa* infection assay

Employing a scratch and infection assay [23], we sought to determine whether or not any of the compounds had the ability to reduce the extent of HCE-T cell lysis caused by endogenous ExoU secreted from the PA103 strain of *P. aeruginosa* (Figure 6). Six hours after scratched HCE-T cells were infected with PA103 (MOI 10), Live/Dead fluorescence microscopy showed that there was extensive cell death along the scratch boarder, with fewer visible detectable live cells, as opposed to no infection, where all cells remained viable and wound closure had begun to occur (Figure 6A). At 1 μM both compounds C and D caused a reduction in the number of cells undergoing lysis at the scratch boarder (Figure 6A), which was accompanied by a decrease in the total amounts of LDH released (Figure 6B). In the presence of both compounds C and D there were fewer lysed cells observed at the scratch boarder and less LDH release, when compared to either compound alone (Figs 6A, B).

**Figure 6:**
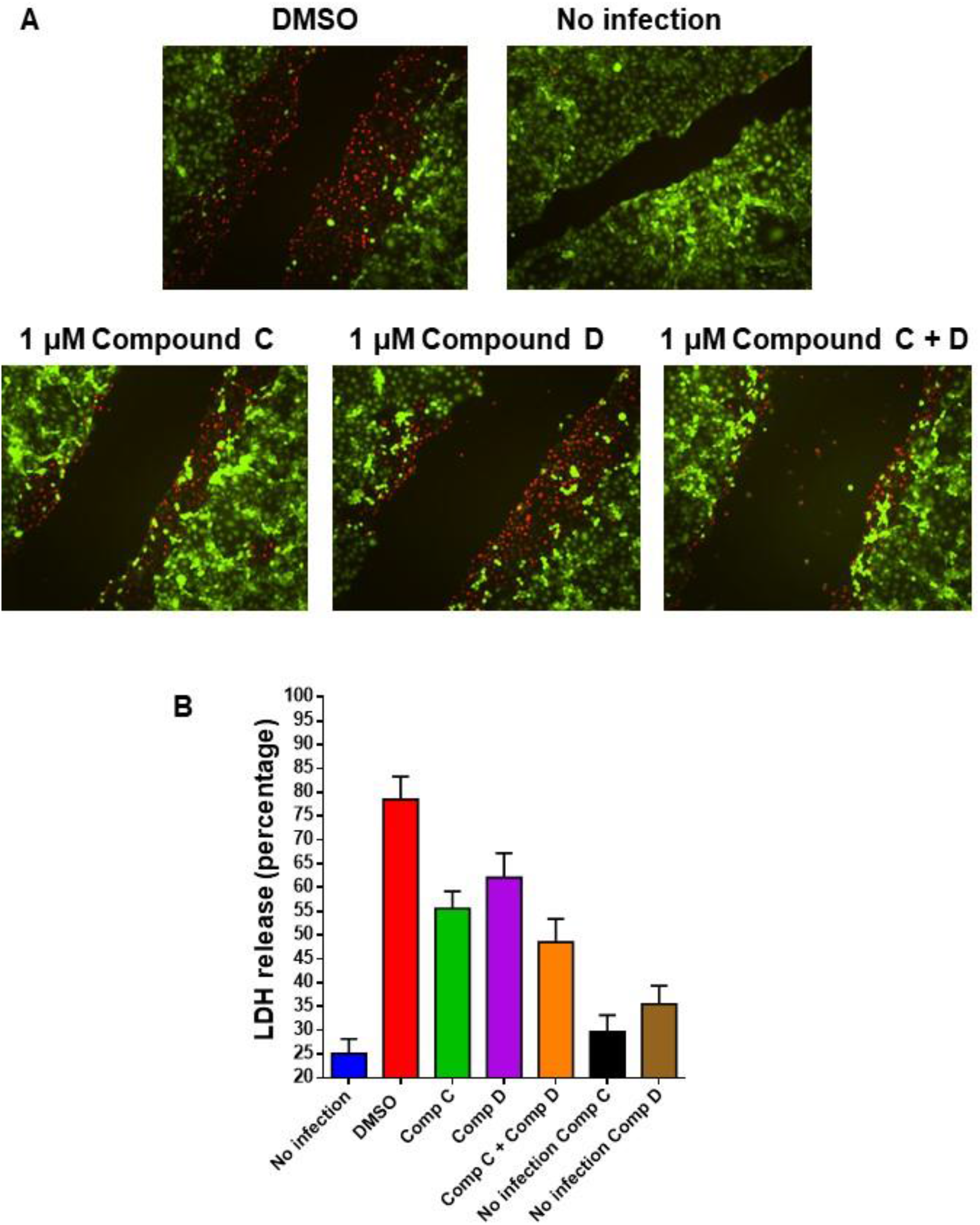
Compounds reduce cell lysis of scratched HCE-T cells after PA103 infection. (A) Live/Dead fluorescence microscopy analysis of HCE-T cells 6 hours after infection with PA103 (MOI 10) in the presence of 1 μM compound C or D, or compounds C and D in combination. (B) LDH assay analysis to quantify cell lysis of HCE-T cells after infection in the presence of 1 μM compound C and D.

## Discussion

In the present study, we employed a high throughput phospholipase assay screen to identify small molecules that have the ability to inhibit recombinant ExoU *in vitro.* Inhibitors of ExoU activity may provide a means by which to interrogate the mechanisms of ExoU activation as well as avenues for potential therapies, in *P. aeruginosa* infections were ExoU is a determinant of poor clinical outcome. Of 6,500 compounds screened, we discovered ~25 compounds that could inhibit ExoU at micromolar concentrations (Figure 1). From this panel we selected compounds C, D, E and F for further analysis.

### Mechanisms of ExoU inhibition

Aside from modulating catalytic activity by binding to the active site, compounds may possess alternative modes of ExoU inhibition [21], for instance by preventing oligomerisation, ubiquitin or phospholipid binding. ExoU oligomerisation in the presence of phospholipids is an important step in its full activation in the host cell. In this study, we showed that of the panel of compounds selected, compound D was able to prevent recombinant ExoU oligomerisation (Figure 2). This distinct mechanism of ExoU inhibition suggested that in combination with other compounds, including compound C synergistic inhibition of ExoU inhibition could be achieved. This was indeed the case as together compounds C and D had a 310 nM IC_50_ (Figure 3). As compound D prevented ExoU oligomerisation, it could be hypothesised that that this molecule disrupts interaction with PIP_2_, perhaps by binding to the ExoU C-terminus. However, the exact binding sites of these compounds on ExoU is not known, therefore, crystal structures and mutagenesis studies will be pivotal in elucidating the modes and mechanisms of compound interaction.

Once inside the host cytoplasm, ExoU binds to ubiquitin and uses its C-terminal 4-helical bundle to localise with and insert into plasma membranes [17]. Consistent with this mechanism, using fluorescence microscopy we observed that GFP-tagged S142A ExoU was localised almost entirely to the plasma membrane. However, in the presence of compound C, much less GFP-ExoU could be detected, and the little ExoU present was found to be colocalised with lysosomes. This result suggests that the binding of compound C to ExoU might cause ExoU to traffic to lysosomes where it is degraded. The human chaperone protein DNAJC5 has recently been elucidated to be indispensable for ExoU cytotoxicity through its trafficking activity, rather than direct chaperoning [22]. Compound C might interfere with ExoU trafficking, or induce an ExoU conformation that promotes its trafficking to lysosomes.

### Therapeutic potential of ExoU inhibitors

Compounds C, D, E and F, as well as inhibiting ExoU phospholipase activity *in vitro* could also mitigate ExoU induced cell lysis in transfected HeLa cells, which suggests potential for therapeutic use. To further explore this, we sought to determine whether or not these compounds had the ability to protect human corneal cells from ExoU secreted by *P. aeruginosa*. Compounds C and D reduced the extent and severity of HCE-T cell lysis after PA103 infection and were more efficacious when used in combination (Figure 6). A combinational therapeutic approach that target distinct mechanisms of ExoU activation might be an attractive strategy. Having nanomolar potency, when compound C and D are used in combination, lends credence to the prospect that they might be efficacious in *in vivo* studies.

### Concluding remarks

ExoU is characterised as the major virulence factor responsible for acute epithelial injury in numerous diseases. Therefore, inhibition of ExoU with small molecules could be an important novel strategy in combating acutely cytotoxic ExoU expressing *P. aeruginosa* infections. Targeting of ExoU may also have the advantage of a decreased risk for selecting resistance, as inhibition of ExoU mediated virulence may synergise with and allow the host immune system to adequately respond [21]. Pharmacological targeting of ExoU may be compatible with antibiotic usage, whereby inhibitors of ExoU serve as an adjuvant therapy. In this way, ExoU inhibitors could mitigate the acute cytotoxic effects whilst conventional antibiotics eliminate the *P. aeruginosa.* ExoU inhibitors, discovered from *in vitro* screening, demonstrate distinct mechanisms of inhibition. These inhibitors should be further investigated, using biochemical and cellular based assays, to advance our understanding of the currently poorly characterised mechanisms of ExoU activity as well as efficacy testing in *in vivo* model disease systems.

